# False-negative assessment of NanoNewron’s NewroBus blood–brain barrier delivery platform in the JAX apical-domain-humanized TfR1 mouse

**DOI:** 10.64898/2026.11.17.767331

**Authors:** Ishita Chakraborty, Tao Yin, Luciano D’Adamio

## Abstract

Humanized transferrin receptor 1 (TfR1) mice are frequently used to evaluate blood–brain barrier (BBB) delivery, yet available strains differ in the extent and design of receptor humanization. We compared two NanoNewron NewroBus-based anti-TNFα constructs, NN-841 (D3 TfR1-binding module) and NN-843 (A2 module), in JAX #038212 (humanized apical domain), Cyagen C001584 (human extracellular-domain CDS model), Cyagen C001860 (human extracellular-domain genomic replacement model), and wild-type controls. Thirty mice received intravenous injections and were evaluated 24 h later. Brain parenchymal-fraction signals were reported in all 12 Cyagen mice, but in none of six JAX or 12 wild-type animals. Serum concentrations were measurable in every group and followed the same rank order for both constructs: Cyagen-CDS > Cyagen-HU > wild type > JAX. Mean CDS-to-JAX serum ratios were approximately 13-fold for both constructs. Compared with the Cyagen-HU strain, the CDS strain showed higher parenchymal concentrations; the ratio of mean parenchymal to mean serum concentration increased prominently for NN-841, but only modestly for NN-843. These findings indicate that model choice influences 24-h circulating concentrations and apparent BBB delivery. Circulating soluble human TfR1-mediated stabilization and endothelial TfR1-dependent transport are plausible, untested explanations for strain-dependent differences. A negative brain-delivery result in JAX #038212 alone cannot be taken as evidence that NewroBus lacks BBB delivery activity

## Introduction

TfR1 is widely exploited for receptor-mediated delivery of biologics across the BBB (1, 2). Because TfR1-targeting antibodies and nanobodies frequently exhibit species-restricted binding, their preclinical evaluation requires animal models expressing a compatible human TfR1 receptor. However, differences in the extent and strategy of receptor humanization may critically influence the suitability of these models for evaluating individual BBB shuttles.

NewroBus is NanoNewron’s platform of engineered humanized TfR1-binding VHH nanobodies developed to deliver therapeutic payloads into the central nervous system (CNS). The platform includes two independently identified families of TfR1-binding nanobodies, A and D, whose members can be incorporated into multifunctional therapeutic constructs (3). We previously demonstrated NewroBus-mediated BBB delivery in genetically humanized TfR1 rats (3, 4). In the present study, we evaluated two anti-TNFα fusion constructs, NN-841 and NN-843, each incorporating two copies of NanoNewron’s humanized anti-TNFα nanobody NN-224 (5), combined with either the D3 (NN-103) or A2 (NN-104) NewroBus module, respectively. Their BBB delivery was compared across three commercially available humanized TfR1 mouse models.

We compared three genetically distinct humanized TfR1 mouse strains. JAX #038212 (C57BL/6-Tfrc tm1(TFRC)Bdes/J) replaces mouse Tfrc residues 196–381 with human TFRC residues 194–379, yielding an apical-domain-humanized receptor. Cyagen C001584 (B6-hTFRC CDS) uses a chimeric coding-sequence strategy to humanize the extracellular domain, while C001860 (B6-huTFRC) uses human genomic DNA to replace the corresponding extracellular-domain region. We asked whether these models would yield concordant measurements of NN-841 and NN-843 entry into brain parenchyma after intravenous administration.

Earlier NewroBus studies in humanized TfR1 rats also revealed differences in systemic exposure between animals with and without a human TfR1 target (3, 4). These findings raise the possibility that receptor humanization affects not only transport at the BBB but also circulating NewroBus concentrations. We therefore examined both serum and brain-fraction measurements in the present mouse comparison.Results

### Extent and strategy of TfR1 humanization differ among the tested mouse strains

The JAX model carries a comparatively limited human apical-domain segment, whereas Cyagen models encode human extracellular TfR1 sequences. The Cyagen models differ in genomic engineering strategy (CDS-based versus genomic replacement). This distinction is illustrated in Figure 1 and prevents direct assumptions of equivalent ligand recognition or transport among the three strains.

**Fig. 1.**
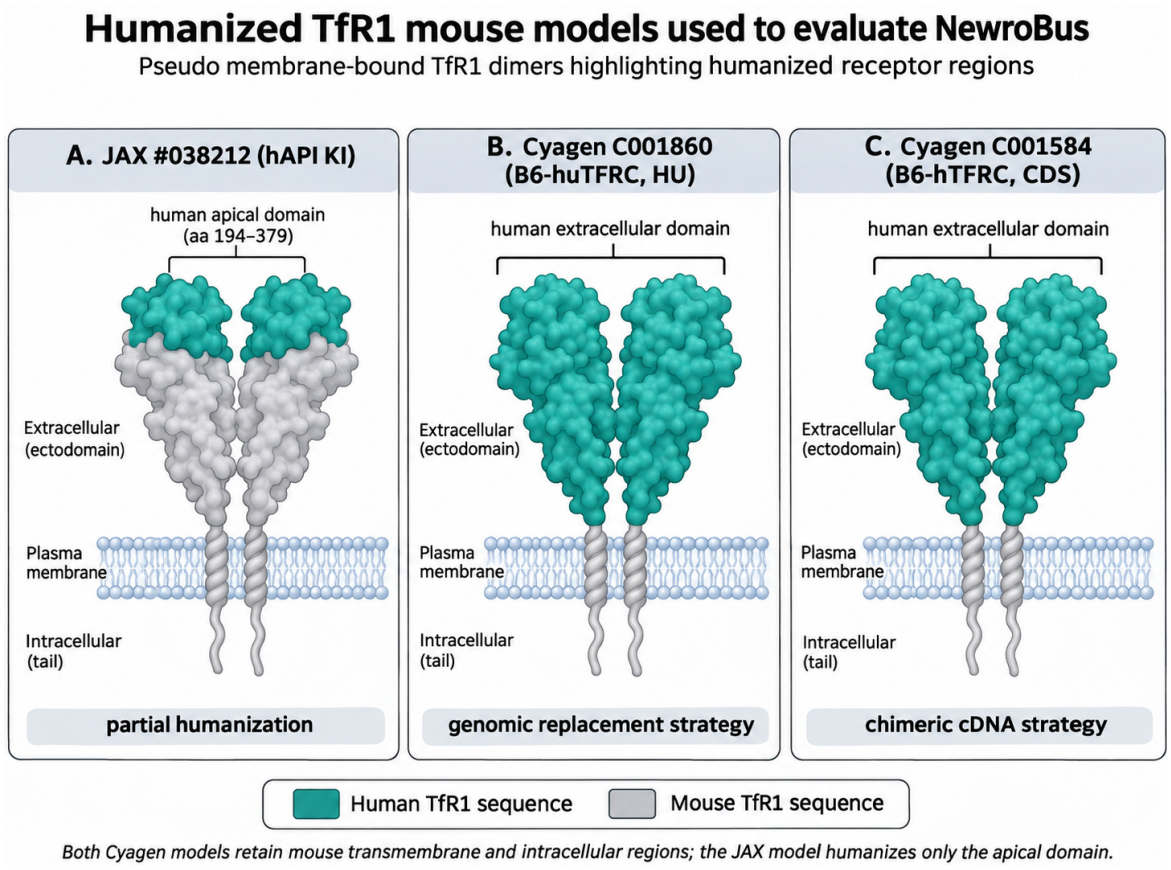
Schematic pseudo-membrane-bound TfR1 dimers illustrating the human-derived (teal) and mouse-derived (gray) receptor regions in JAX #038212, Cyagen C001860 (HU), and Cyagen C001584 (CDS). Structural shapes are conceptual and not atomistic models.

### NewroBus exposure and brain delivery differ by mouse strain

Thirty mice were assigned to eight construct-by-genotype groups (15 mice per construct): six wild-type, three JAX, three Cyagen C001584 (CDS), and three Cyagen C001860 (HU) mice for each construct. They were injected intravenously and collected 24 h later. Individual animal labels, sex, age, weight, calculated dosing volume, and formulation concentrations are tabulated in Table 1.

**Table 1.** Characteristics and dosing information for individual mice used in the NewroBus biodistribution study. The table lists animal identification numbers, TfR1 genotype/model, sex, age at injection (days), body weight (g), administered nanobody, intravenous (IV) injection volume, and formulation concentrations. Mice received either NN-841 (D3 NewroBus) or NN-843 (A2 NewroBus) at an injection volume of 1 µL/g body weight. Four mouse groups were examined: wild-type (WT), JAX #038212, Cyagen C001860 (HU), and Cyagen C001584 (CDS). Each nanobody was administered to 15 mice (WT, n = 6; JAX, n = 3; HU, n = 3; CDS, n = 3), for a total of 30 animals. Blood and brain tissues were collected 24 hours after injection. CSF was not collected.

| Animal | Sex | Age (d) | Wt (g) | Construct | IV vol (µL) | Molarity of dosing solution (µM) |
|---|---|---|---|---|---|---|
| **WT** | M | 202 | 29 | NN841 | 29.0 | 32.855 |
| **WT** | F | 231 | 24 | NN841 | 24.0 | 32.855 |
| **WT** | F | 208 | 22.5 | NN841 | 22.5 | 32.855 |
| **WT** | F | 208 | 24 | NN841 | 24.0 | 32.855 |
| **WT** | M | 202 | 28 | NN841 | 28.0 | 32.855 |
| **WT** | M | 99 | 23 | NN841 | 23.0 | 32.855 |
| **Jax** | M | 190 | 34 | NN841 | 34.0 | 32.855 |
| **Jax** | F | 190 | 26 | NN841 | 26.0 | 32.855 |
| **Jax** | F | 190 | 29 | NN841 | 29.0 | 32.855 |
| **Cy-HU** | M | 133 | 35 | NN841 | 35.0 | 32.855 |
| **Cy-HU** | M | 133 | 33.2 | NN841 | 33.2 | 32.855 |
| **Cy-HU** | M | 417 | 17.3 | NN841 | 17.3 | 32.855 |
| **Cy-CDS** | F | 133 | 23.2 | NN841 | 23.2 | 32.855 |
| **Cy-CDS** | F | 133 | 24.3 | NN841 | 24.3 | 32.855 |
| **Cy-CDS** | F | 133 | 22.3 | NN841 | 22.3 | 32.855 |
| **WT** | M | 202 | 37 | NN843 | 37.0 | 35.4182 |
| **WT** | F | 209 | 22.6 | NN843 | 22.6 | 35.4182 |
| **WT** | F | 209 | 25 | NN843 | 25.0 | 35.4182 |
| **WT** | M | 232 | 27 | NN843 | 27.0 | 35.4182 |
| **WT** | M | 202 | 28 | NN843 | 28.0 | 35.4182 |
| **WT** | M | 202 | 29 | NN843 | 29.0 | 35.4182 |
| **Jax** | M | 190 | 30 | NN843 | 30.0 | 35.4182 |
| **Jax** | F | 190 | 29 | NN843 | 29.0 | 35.4182 |
| **Jax** | F | 190 | 28 | NN843 | 28.0 | 35.4182 |
| **Cy-HU** | F | 134 | 27 | NN843 | 27.0 | 35.4182 |
| **Cy-HU** | F | 134 | 27 | NN843 | 27.0 | 35.4182 |
| **Cy-HU** | F | 134 | 27 | NN843 | 27.0 | 35.4182 |
| **Cy-CDS** | F | 134 | 21 | NN843 | 21.0 | 35.4182 |
| **Cy-CDS** | F | 134 | 32 | NN843 | 32.0 | 35.4182 |
| **Cy-CDS** | F | 134 | 39 | NN843 | 39.0 | 35.4182 |

Serum concentrations of NN-841 and NN-843 (Figure 2) were detectable in all mouse groups, with the highest levels observed in Cyagen-CDS mice, followed by Cyagen-HU, WT, and JAX mice. Mean serum concentrations at 24 h were 171.6, 88.2, 623.0, and 1140.2 pM for NN-841 in WT, JAX, Cyagen-HU, and Cyagen-CDS mice, respectively; the corresponding NN-843 concentrations were 254.5, 108.5, 1045.5, and 1512.3 pM (Table 1; Figure 2). The CDS/JAX ratios of mean serum concentrations were 12.9 for NN-841 and 13.9 for NN-843, whereas the CDS/HU ratios were 1.83 and 1.45, respectively.

**Fig. 2.**
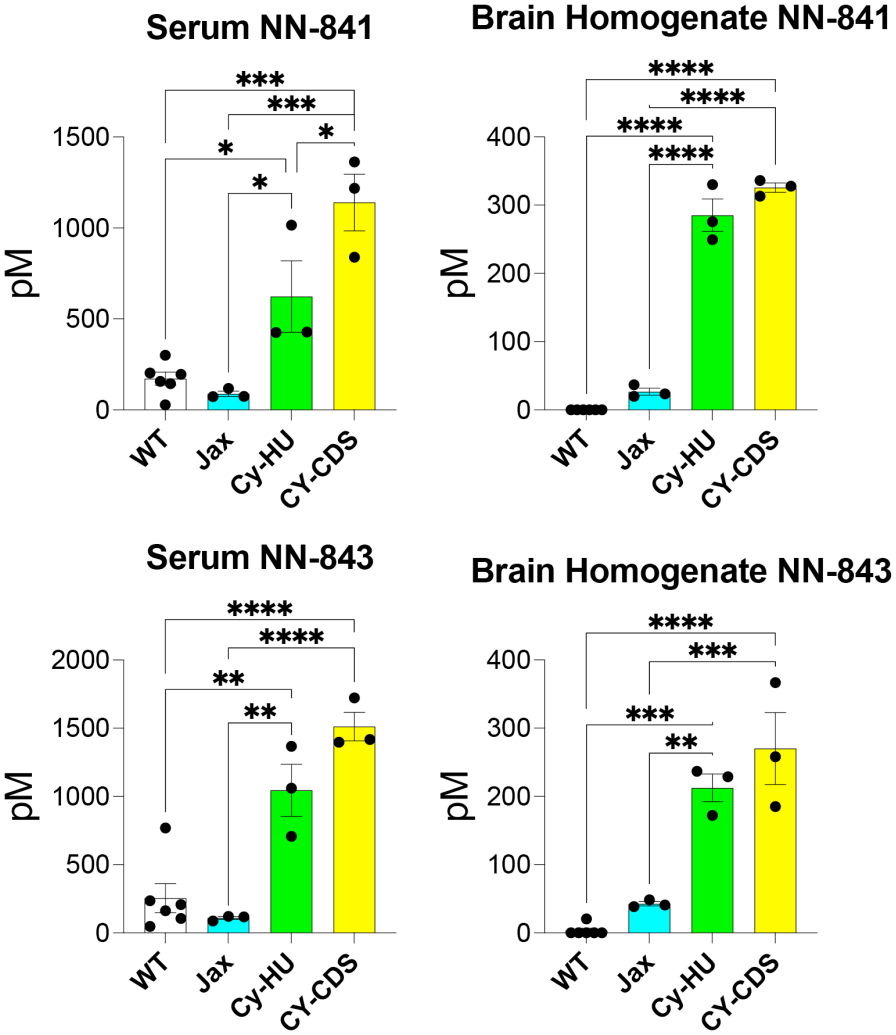
NN-841 and NN-843 concentrations in serum and unfractionated brain homogenate (raw) 24 h after IV administration. Individual animal values, means and error bars are retained from Prism. Statistical analyses were performed using GraphPad Prism. Differences among experimental groups were evaluated by ordinary one-way analysis of variance (ANOVA), followed by Tukey’s multiple-comparisons test to assess pairwise differences between groups. Adjusted P values < 0.05 were considered statistically significant. Statistical significance is indicated as follows: *P < 0.05, **P < 0.01, ***P < 0.001, and ****P < 0.0001. ns, not significant (P ≥ 0.05).

Whole-brain raw nanobody concentrations showed marked differences among mouse models. Mean concentrations of NN-841 were 0, 26.6, 285.3, and 325.7 pM in WT, JAX, Cyagen-HU, and Cyagen-CDS mice, respectively; corresponding NN-843 concentrations were 3.4, 42.6, 212.6, and 270.1 pM. Compared with JAX mice, whole-brain concentrations in Cyagen-CDS mice were 12.2-fold higher for NN-841 and 6.3-fold higher for NN-843, whereas those in Cyagen-HU mice were 10.7- and 5.0-fold higher, respectively. Thus, both NewroBus constructs exhibited substantial brain accumulation in the Cyagen models, whereas accumulation was negligible in WT mice and markedly reduced in JAX mice.

### Model-dependent accumulation of NewroBus in brain vascular and parenchymal fractions

To further characterize brain distribution, whole-brain homogenates, vascular-enriched fractions, and capillary-depleted parenchymal fractions were analyzed after normalization to their respective protein concentrations (Figure 3). For NN-841, normalized whole-brain concentrations were markedly higher in Cyagen-HU and Cyagen-CDS mice than in WT and JAX mice. Mean vascular concentrations were 2.01 and 9.82 pM/(mg/mL) protein in Cyagen-HU and Cyagen-CDS mice, respectively, representing a 4.9-fold difference. The corresponding parenchymal concentrations were 22.16 and 106.83 pM/(mg/mL) protein, a 4.8-fold difference. Similarly, NN-843 accumulated predominantly in the Cyagen models, with mean vascular concentrations of 2.08 and 5.88 pM/(mg/mL) protein in Cyagen-HU and Cyagen-CDS mice, respectively, and corresponding parenchymal concentrations of 56.91 and 83.82 pM/(mg/mL) protein. In contrast, vascular and parenchymal concentrations of both nanobodies were undetectable in WT and JAX mice. These results demonstrate substantial NewroBus accumulation in the capillary-depleted brain parenchymal fractions of both Cyagen models, whereas the JAX humanized TfR1 mouse yielded essentially negative results for both constructs.

**Fig. 3.**
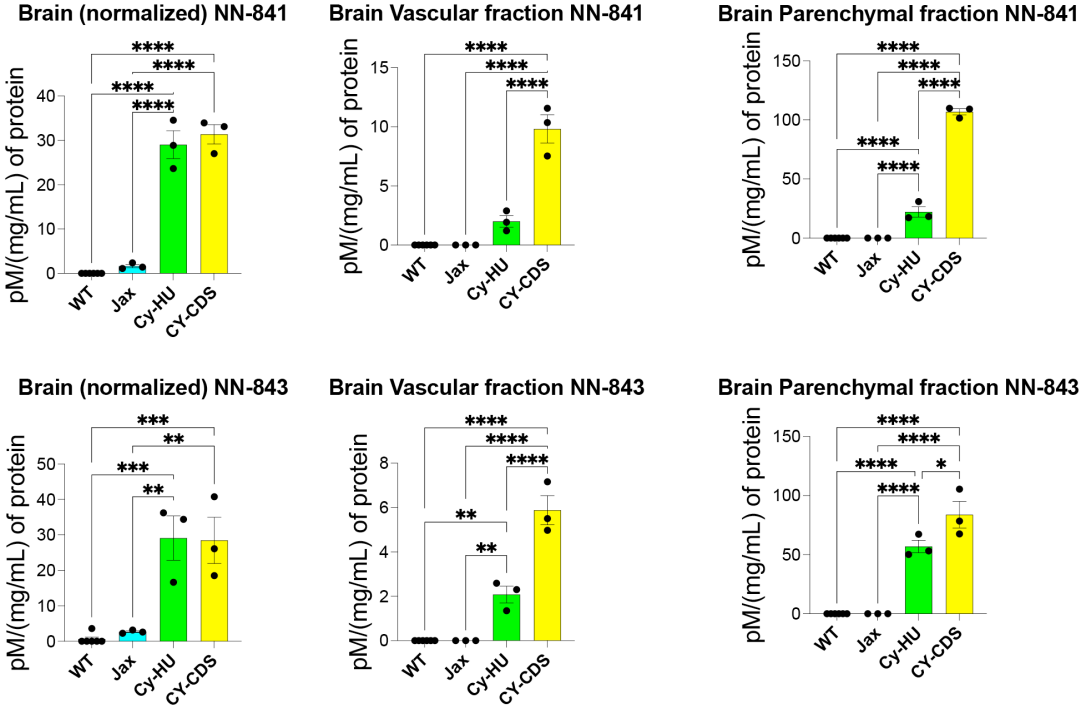
Model-dependent accumulation of NN-841 and NN-843 in whole-brain homogenates, vascular-enriched fractions, and capillary-depleted brain parenchymal fractions. Whole-brain homogenates analyzed in Fig. 2 were fractionated into vascular-enriched and capillary-depleted parenchymal fractions. Nanobody concentrations were normalized to the protein concentration of each sample, determined by Bradford assay, and expressed as pM/(mg/mL) of protein. Each dot represents an individual animal; bars indicate mean ± SEM (WT, n = 6; Jax, Cy-HU, and Cy-CDS, n = 3 per construct). Statistical comparisons were performed using ordinary one-way ANOVA followed by Tukey’s multiple-comparisons test. **P < 0.01, ***P < 0.001, ****P < 0.0001.

Relative to both Cyagen models, JAX #038212 gave a negative parenchymal-fraction result for each NewroBus-containing construct, although the same compounds were detected in serum. The data demonstrate that the JAX hAPI mouse is not an appropriate stand-alone negative test for NN-841 or NN-843 brain delivery. They do not establish the molecular explanation for this incompatibility, whether the receptor binds the two constructs in situ, or whether other TfR1 shuttles behave similarly. Critically, an independently conducted evaluation based solely on JAX #038212 could mistakenly assign the negative outcome to the NewroBus platform rather than model choice.

## Discussion

Our findings demonstrate that the choice of humanized TfR1 mouse model profoundly influences the apparent brain delivery of NewroBus constructs. Although both NN-841 and NN-843 accumulated substantially in the brain parenchymal fractions of Cyagen-HU and Cyagen-CDS mice, neither construct was detectable in the corresponding fractions of JAX #038212 mice. Thus, the JAX model yielded results comparable to wild-type mice, despite expressing a partially humanized TfR1 receptor.

An additional observation was the marked difference in circulating NewroBus concentrations among the mouse models. Serum concentrations of both constructs were lowest in JAX mice, like those observed in wild-type animals, and substantially higher in both Cyagen models, particularly Cyagen-CDS. This pattern is consistent with our previous findings in genetically humanized TfR1 rats, in which expression of human TfR1 was associated with increased circulating concentrations of NewroBus constructs (1,2). Together, these observations suggest that interactions with human TfR1 may contribute not only to brain delivery but also to systemic persistence.

One possible mechanism involves binding to circulating soluble human TfR1 (sTfR1), which could increase the effective molecular size of NewroBus constructs and reduce their renal clearance. Alternatively, differences in membrane-bound TfR1 expression, receptor-mediated disposition, or soluble receptor abundance among the models could contribute to the observed serum concentrations. The particularly high serum and brain concentrations in Cyagen-CDS mice may reflect greater availability of functional human TfR1, although receptor abundance was not directly measured. These possibilities remain speculative, and the present experiments do not establish the mechanism underlying the pharmacokinetic differences.

Importantly, the low serum concentrations in JAX mice could themselves contribute to the absence of detectable brain parenchymal accumulation, independently of any differences in TfR1 binding or transcytosis. Thus, the negative brain-delivery results could reflect inadequate receptor recognition, reduced systemic persistence, inefficient receptor-mediated transport, or a combination of these factors. Distinguishing among these possibilities is not necessary to establish the central conclusion of this study.

Regardless of the underlying mechanism, JAX #038212 does not provide a suitable experimental model for evaluating NewroBus-mediated brain delivery under the conditions tested. The same two constructs that yielded essentially negative results in JAX mice demonstrated substantial accumulation in the capillary-depleted brain parenchymal fractions of both Cyagen humanized models. Consequently, independent evaluation of NewroBus using JAX #038212 could generate false-negative conclusions regarding the platform’s BBB delivery capability. These findings underscore that humanization of TfR1 does not automatically establish compatibility with a particular BBB shuttle and that negative results obtained in an unvalidated humanized animal model should not be interpreted as evidence of therapeutic delivery failure.

## Materials and methods

### Mouse models and experimental design

Animals were wild-type mice, JAX strain #038212 (C57BL/6-Tfrc tm1(TFRC)Bdes/J), Cyagen C001584 (B6-hTFRC CDS), or Cyagen C001860 (B6-huTFRC). Both male and female mice were studied; age, weight, sex, individual animal labels, and administration dates are given in Table S1. Nanobodies were injected IV on April 7, 12, or 13, 2026, and specimens were harvested 24 h after dosing. Procedures for nanobody injection, blood collection, brain isolation, capillary enrichment/depletion, fraction processing, and nanobody assays followed those previously reported for the humanized TfR1rat studies (3, 4), with the exception that cerebrospinal fluid was not collected from mice because collection was not feasible.

### NewroBus constructs and administration

NN-841 and NN-843 are anti-TNFα fusion nanobodies containing the D3 and A2 NewroBus modules, respectively. Both constructs were administered intravenously at 1 µL/g body weight (1 mL/kg) with stock concentrations of 32.855 µM for NN-841 and 35.4182 µM for NN-843, equivalent to nominal doses of 32.855 and 35.4182 nmol/kg, respectively.

### Tissue collection and fractionation

All mouse procedures followed the NewroBus humanized-rat experimental procedures previously described (3, 4), except that cerebrospinal fluid (CSF) could not be harvested from these mice. At the 24-h terminal collection, blood was obtained for serum preparation, and mice were perfused before brain removal, following the prior rat protocol. The previous protocol used transcardiac perfusion with cold calcium- and magnesium-free PBS (10 mL/min for 10 min). Choroid plexuses were removed and brain tissue was homogenized on ice in S1 buffer (250 mM sucrose, 20 mM Tris, pH 7.4, 1 mM EDTA and 1 mM EGTA) using a glass–Teflon homogenizer. Following centrifugation at 1,000 × g for 10 min at 4 °C, pellets were resuspended in 17% (w/v) 60-kDa dextran and centrifuged at 4,200 × g for 15 min. The resulting pellet constituted the vascular-enriched P1 fraction. The supernatant was diluted in S1 buffer and centrifuged again at 4,200 × g for 15 min to obtain the vascular-depleted parenchymal PC fraction. Fractions were lysed in inhibitor-containing buffer and sonicated. The concentrations of NN-841 and NN-843 were determined with the previously described nanobody immunoassay method (3, 4). Specific dilution factors, quantitative assay thresholds and specimen recovery were not available in the present source spreadsheet and should be confirmed against the experimental records.

### Brain protein determination and concentration normalization

Nanobody concentrations measured in whole-brain homogenates, brain vascular fractions, and brain parenchymal fractions were normalized to the total protein concentration of the corresponding samples, determined using the Bradford assay. Normalized values were calculated by dividing the measured nanobody concentration (pM) by the protein concentration (mg/mL) and are expressed as pM/(mg/mL).

### Data analysis and interpretation

The mouse was the experimental unit. Concentrations are summarized as group means ± SEM. Comparisons across the four mouse groups were performed using ordinary one-way analysis of variance (ANOVA), followed by Tukey’s multiple-comparisons test, as specified for the supplied GraphPad Prism analyses. Adjusted P values < 0.05 were considered significant; *P < 0.05, **P < 0.01, ***P < 0.001, and ****P < 0.0001. For descriptive cross-model comparisons, ratios were calculated from group means rather than from within-animal paired ratios. Because all measurements were obtained at a single 24-h time point, serum area under the concentration–time curve and intrinsic BBB transport rates could not be determined.

